# Advancing Transcription Factor Binding Site Prediction Using DNA Breathing Dynamics and Sequence Transformers via Cross Attention

**DOI:** 10.1101/2024.01.16.575935

**Authors:** Anowarul Kabir, Manish Bhattarai, Kim Ø. Rasmussen, Amarda Shehu, Alan R Bishop, Boian Alexandrov, Anny Usheva

## Abstract

Understanding the impact of genomic variants on transcription factor binding and gene regulation remains a key area of research, with implications for unraveling the complex mechanisms underlying various functional effects. Our study delves into the role of DNA’s biophysical properties, including thermodynamic stability, shape, and flexibility in transcription factor (TF) binding. We developed a multi-modal deep learning model integrating these properties with DNA sequence data. Trained on ChIP-Seq (chromatin immunoprecipitation sequencing) data *in vivo* involving 690 TF-DNA binding events in human genome, our model significantly improves prediction performance in over 660 binding events, with up to 9.6% increase in AUROC metric compared to the baseline model when using no DNA biophysical properties explicitly. Further, we expanded our analysis to *in vitro* high-throughput Systematic Evolution of Ligands by Exponential enrichment (SELEX) and Protein Binding Microarray (PBM) datasets, comparing our model with established frameworks. The inclusion of DNA breathing features consistently improved TF binding predictions across different cell lines in these datasets. Notably, for complex ChIP-Seq datasets, integrating DNABERT2 with a cross-attention mechanism provided greater predictive capabilities and insights into the mechanisms of disease-related non-coding variants found in genome-wide association studies. This work highlights the importance of DNA biophysical characteristics in TF binding and the effectiveness of multi-modal deep learning models in gene regulation studies.

## Introduction

Knowing the sequence specificities of DNA- and RNA-binding proteins, also known as transcription factors (TFs), is essential for understanding the regulatory processes in biological systems and for identifying causal disease variants. These processes are attributed to the fact of regulating many biological activities, such as transcription, translation, or suppression, by influencing gene level expression. Understanding the impact of these variants on TF binding and gene regulation remains a key area of research, with implications for unraveling the complex mechanisms underlying various genetic disorders. Much of this interest are due to the over 98% of the human genome is non-coding, the function of which is not very well-defined. A model that can predict function directly from sequence may reveal novel insights about these non-coding elements. Over 1200 genome-wide association studies have identified nearly 6500 disease- or trait-predisposing single-nucleotide polymorphisms (SNPs), 93% of which are located in non-coding regions (1), highlighting the importance of such a predictive model.

Building upon the reality of transcription and TF binding site prediction, it is crucial to recognize that the active transcription initiation and assistance are significantly influenced by the binding of TFs to DNA. This complex interaction is modulated by a myriad of epigenomic mechanisms, notably the local biophysical properties of DNA. Critical factors such as local thermodynamic stability and flexibility play a pivotal role in TF-DNA binding (2; 3; 4). The structural integrity of DNA, largely dependent on hydrogen bonds, is a key component in this interaction. These hydrogen bonds, being weaker than covalent bonds, render DNA susceptible to conformational changes due to thermal fluctuations at physiological temperatures. This phenomenon, known as “DNA breathing dynamics” or “DNA bubbles,” involves the spontaneous local opening and closing of the DNA double helix, significantly impacting the flexibility of DNA at specific sites (5; 6; 7; 8; 9; 10; 11; 12; 13; 14; 15; 16; 17). These dynamic movements of DNA strands have profound implications for TF binding and gene expression, as evidenced by numerous studies. The propensity of DNA to form these “bubbles” and the features associated with them, such as base-pair flipping, bubble length, and height, are vital for understanding TF binding. Intriguingly, many of these dynamic features are influenced by nucleotides that lie outside the TF motifs. This insight offers a window into the mechanisms through which functional non-coding variants, often implicated in disease. We utilize these physics-based biophysical properties explicitly in this study to guide the state-of-the-art model to understand TF-DNA binding sites.

There has been a growing interest to predict function directly from sequence, instead of using curated datasets such as gene models and multiple species alignment. Based on the nature of the high-throughput technologies, the experimental datasets for TF-DNA binding comes in different qualitative and quantitative formats. Protein Binding Microarray (PBMs) (18; 19) provide a specificity coefficient for each probe sequence. High throughput Systematic Evolution of Ligands by Exponential enrichment (SELEX) (HT-SELEX) (20) generates a set of very high affinity sequences. These datasets comes in large quantity, typically high-throughput experiment measures between 10, 000 and 100, 000 sequences. On the other hand, *in vivo* Chromatin immunoprecipitation sequencing (ChIP-seq) provides a ranked list of putatively bound sequences of varying length. ChIP-seq reads often localize to “hyper-ChIPable” regions of the genome near highly expressed genes. Different line of works have been proposed to mitigate these challenges. However, existing approaches that are applicable to both *in vivo* and *in vitro* experiments are limited.

There has been a variety of deep learning based methods for automatically predict nature of these non-coding regions, specifically TF-DNA binding regions. Previous works mostly depend solely on the Convolutional neural network (CNN) architectures. Generic reasons for that is the capability of CNNs to extract local and hierarchical features from the given input, which is a major condition for finding the binding motifs in the genome. The weight sharing capacity with different size of kernels and many filters provide major boost in performance of this area of work. In the earlier day DeepBind (21) and DeepSEA (22) started utilizing CNNs and their evaluation beats the non deep learning based methods. These two works mostly varies in the setting in the problem and usage of the datasets. Specifically, DeepSEA (22) applies multi-tasks learning where the tasks are TF binding prediction by predicting chromatin features with single-nucleotide sensitivity, DNase I–hypersensitive sites and Histone marks profiling. As such the dataset was ChIP-seq experiment based chromatin features prediction genome-wide. DeepBind (21), on the other hand, applies the problem in single-tasks learning, in diverse set of experimental data both *in vitro* and *in vivo* set. Later DeeperBind (23) was proposed as an extension over DeepBind which uses Long Short-Term Memory (LSTM) network on top of CNNs. There are some other solely CNN based architecture that studies the problem that trained on the *in vitro* SELEX datasets and show the applicability on both *in vivo* and *vitro* settings. For instance, WSCNN (24) uses a weakly supervised CNN architecture and DeepSELEX (25) uses 963 in vitro experiments corresponding to 32 TFs. The CNNs in the HOCNN (26) is high-order and tested on to 214 ChIP-seq experiments across five cell lines.

As DeeperBind (23) utilizes CNNs followed by LSTM architecture, an immediate improvement in the architecture of adding biLSTM in DanQ (1) instead of LSTM showed models applicability in the human genome-wide TF-DNA binding prediction on the same datasets of predicting 919 chromatin features from DeepSEA (22). Later by understanding the broad applicability of trying to enhance the interpretability of the deep learning models TBiNet (27) and DeepGRN (28) introduce attention (29) in their model. On the differential side of these two models, TBiNET architecture consists of CNNS followed by the attention component and then applying a bidirectional LSTM module. The intention was to able to explain the CNN kernels in terms of motif discovery and to capture the large and long context of the input DNA sequence part of which may or may not have TF binding sites. Contrarily, DeepGRN’s attention is at the end, specifically CNNs, BiLSTM and an attention module. Note that DeepGRN was developed by using the training datasets from the 2016 ENCODE DREAM *in vivo* TF binding prediction Challenger. In this challenge, the models are trained on all chromosomes except 1, 8, and 21, and chromosome 11 is used as validation; while TBiNET, DanQ follows the DeepSEA dataset split, specifically chromosome 8 and 9 for the test set and chromosome 7 as the validation set.

Recently, foundation models particularly on transformer neural networks (29) and its subsequent improvement becomes popular in natural language, protein as well as genomic language domain. DNABERT2 (30) is one among many state-of-the-art works. It is trained on multi-species genome with Byte Pair Encoding (BPE) instead of k-mer tokenization. It generalizes in many epigenomic downstream tasks including TF-DNA binding sites prediction and variant effect estimation. A major benefit of applying these deep learning based models is to exploit the fact of removing the generation of hard-crafted features. Contrarily, no domain knowledge challenges these models with explainability while performing to the great extent. In this study, we utilize both the fact that a genomic language model in a multi-modal setting as a physics guided system to accurately predict TF-DNA binding events in human genome.

Recent research has explored the utilization of various DNA features to enhance TF binding specificity predictions. The studies by (31; 32; 33) have leveraged shape features such as Minor Groove Width (MGW), Roll, Propeller Twist (ProT), and Helix Twist (HelT), which were extracted from Monte Carlo (MC) simulations (34). These investigations have shown that predictions of TF binding specificity are significantly enhanced when both sequence and shape models are employed, as opposed to relying on sequence data alone. Similarly, The DLBSS framework (35) applied deep CNN architecture to improve the performance of predicting *in vitro* PBM datasets with DNA shape features. DNAffinity (36) uses structural and mechanical DNA properties directly derived from atomistic molecular dynamics (MD) simulations. Similar to DLBSS (35) it is trained on *in vitro* datasets, however utilizes random forest regressor for predicting TF-DNA binding affinity. A common ground across these features and frameworks is that they enhance TF binding performances for either *in vivo* or *in vitro* datasets, where our feature aids for the deep leaning framework to enhance the binding performance for both *in vivo* and *in vitro* datasets.

Considering the advances in the TF-DNA binding prediction tasks, we first aim for guiding a foundation model pretrained solely on DNA sequences. Particularly, we leverage DNA breathing dynamics in a cross-attention aggregation module to enhance the model’s predictive accuracy. The intuition of such effective design of the model underlies based on the assumption that the breathing and foundational features might come from different modalities. We rigorously tested our model using diverse datasets both *in vivo* and *in vitro* to validate its robustness and versatility. The evaluation *in vivo* datasets shows that our proposed model improves at least 3% on average in both AUROC and AUPR metrics. More ingrained findings demonstrate that the model significantly improves prediction performance in over 660 binding events out of 690, with up to 9.6% increase in AUROC metric compared to the baseline model when using no DNA biophysical properties explicitly. Furthermore, we allow model, particularly cross-attention weights, to find evolutionary enriched motif regions. This synthesizes that the high-throughput-derived biophysical DNA characteristics with sequence data offers a more nuanced understanding of TF binding sites. Further experiments *in vitro* SELEX and gcPBM datasets shows efficacy of the breathing dynamics over sequence alone information, particularly at least 3% improvements on *R*^2^ scale.

We developed an innovative computational framework for predicting TF binding sites that extends beyond traditional DNA sequence analysis. A key feature of our model is its integration of biophysical characteristics of TFs, which significantly enhances performance during testing. This integration provides a deeper understanding of TF-DNA interactions and contributes to more accurate and comprehensive TF binding site predictions. Crucially, our framework is not limited to a specific TF group, thereby enabling its application to arbitrary TFs, including those not encountered during training. This broad applicability is further bolstered by our model’s transfer learning capabilities, demonstrated through training and testing across various cell types. Such versatility is pivotal in assessing the functional impact of variants, as it allows for the comparison of TF binding intensities between altered sequences and reference sequences. To the summary we list the core contributions of this study towards TF-DNA binding in the following.

- Application of DNA breathing dynamics to guide a foundation model to exploit different feature modalities.
- Integration of cross-attention framework in multi-modal settings.
- Explainability of cross-attention weights for finding motif enriched regions.
- Broad evaluation on TF-DNA binding data *in vivo* and *in vitro*.

Our research aims to deepen the understanding of gene regulation mechanisms and open new avenues for identifying novel therapeutic targets. By bridging the gap between DNA sequence data and the dynamic biophysical properties of DNA, our model represents a significant step forward in the computational prediction of TF binding sites.

## Materials and methods

### TF-DNA binding affinity dataset

TFs are the central regulatory modular proteins that recognize fragments of DNA (typically 6–20 bp long) for RNA polymerizes that trigger the subsequent transcription (36) process. To derive the utility of the bio-physical characteristics generated by the pyDNA-EPBD (37), we analyze the TF-DNA binding affinity prediction tasks both *in vitro* and *in vivo* derived datasets.

### *In vitro* dataset: HT-SELEX and gcPBM

For a deeper understanding of the contribution of the EPBD models, we utilized ground truth binding affinities from two different *in vitro* experimental setup, such as genomic context protein binding microarrays (gcPBM) (31) and high-throughput SELEX experiments (HT-SELEX) (38). These experimental datasets come from different experimental assumptions and as such different numerical forms which makes the computational modelling more challenging to capture the essentiality. For instance HT-SELEX generates high affinity sequences for human transcription factors, and PBMs provide specificity coefficient for each probe sequence (21).

We utilize the pre-processed data described by (39). The authors reported TF-DNA binding specificities for all DNA sequences of length *M*, also known as M-words. This data went through many filtering steps, such as high M-word variability, deep read coverage, selection of core motifs for each TF, and filtering out infrequent M-words. At the end, this set contains specificity information for 215 TFs from 27 families of the HT-SELEX dataset (available in European Nucleotide Archive [ENA]: PRJEB14744, downloaded from^1^ (file name: M-word scores, last accessed: 12 February, 2024 13:28)). The data corresponds to the relative affinity scores for the corresponding M-words. It contains total 1, 788, 827 number of sequences where C2H2-SP3 contains the minimum number of sequences (1, 013) and homeodomain-RHOXF1 contains the maximum number of sequences (62, 871). Figure 1 depicts the affinity distribution across 215 TFs in ascending order by the average value (shown in orange). The green bar depicts the standard deviation and the gray bar demonstrates minimum and maximum value. The distribution of this dataset shows similar variance across different TFs while containing the same maximum value which poses a major challenge for the computational learning model. Note that the sequence length in this dataset varies from 9 to 15 nucleotide base pairs.

**Fig. 1:**
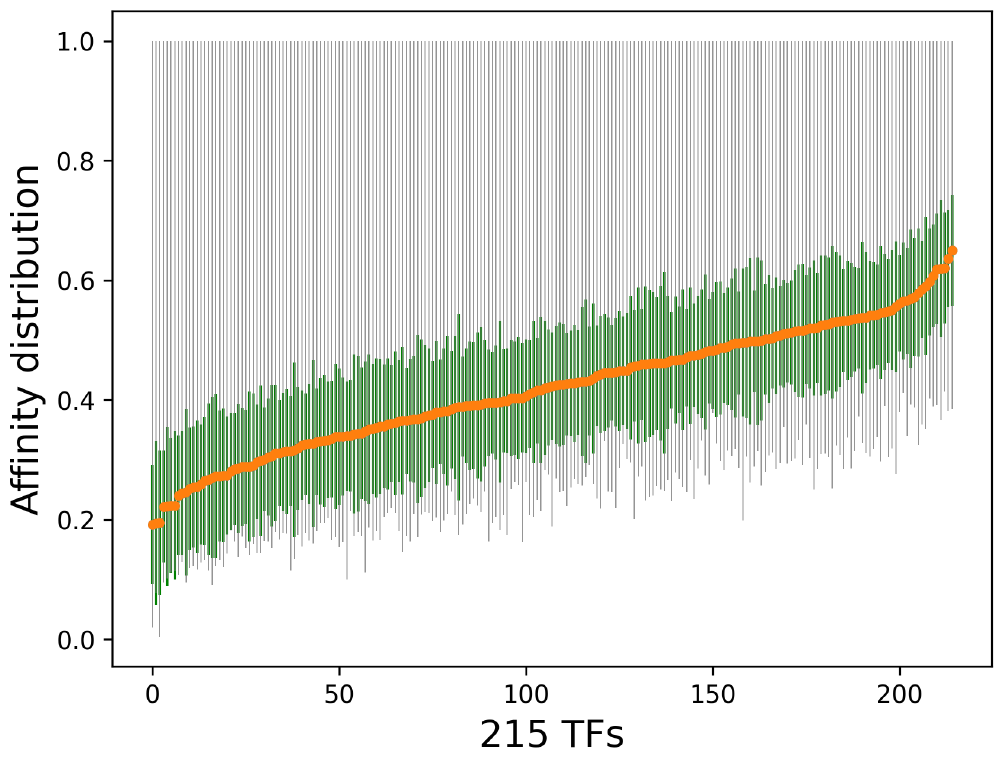
Affinity distribution across 215 TFs in ascending order by the average value (shown in orange). The green bar depicts the standard deviation and the gray bar demonstrates minimum and maximum value. Overall it shows similar variance across different TFs while containing the same maximum value, posing a challange for the computational model.

The gcPBM (18) applies four protein binding microarray (PBM) experiments for binding GST-tagged human transcription factors Mad2(Mxi1), Max, and c-Myc to double-stranded 180K alignment microarrays in order to determine their binding specificity for putative DNA binding sites in native genomic context. The dataset (40) contains 23, 028 sequences where all sequences are of length 36. The specificity value distributions as in mean and standard deviation are 8.650 (±0.801), 4.091 (±0.214) and 8.647 (±0.792) for Mad, Max and Myc, respectively. The varied nature of the experiments and as such the ground-truth values pose extra challenges to apply a single model to demonstrate its applicability in all settings. We studied breathing dynamics generated by pyDNA-EPBD (37) on these datasets and show the efficacy of our physics guided model.

### *In vivo* dataset: Human TF-DNA ChIP-seq experiments

To understand and evaluate the TF-DNA prediction tasks *in vivo* we utilize a comprehensive set of human TF binding sites based on ChIP-seq (chromatin immunoprecipitation sequencing) experiments, where the chromatin is immunoprecipitated by TF-specific antibody and the retained chromatin is then sequenced. The data (regions of enrichment also known as peak calls) we used in this study is generated by the ENCODE (Encyclopedia of DNA Elements) Analysis Working Group (AWG) based on a uniform processing pipeline from September 2007 through the March 2012. The dataset contains 690 ChIP-seq experiments representing 161 unique regulatory factors spanning 91 human cell types. The experiments are conducted in various treatment conditions with two different experiment quality as good or caution. Our study neither discriminate among the treatment conditions nor evaluate separately based on experiments quality.

To prepare the inputs for the labels we follow the DeepSEA (22) protocol thoroughly. To the best of our knowledge DeepSEA dataset is not updated recently which might be incongruous with the recent human genome, as such we generate our own train, validation and test set using the same protocol. The 690 TF-DNA binding experiments are collected from UCSC browser^2^. We also download the human reference genome GRCh37/hg19 from UCSC browser^3^ (file name: hg19.fa.gz, last modified: 2020-01-17 13:58, and file size: 934M). The corresponding chromosome sizes are downloaded from the same URL as of the genome, (file name: hg19.chrom.sizes, last modified: 2020-01-15 16:40, file size: 7.0K). Following DeepSEA (22), DanQ (1) and FactorNet (41), we only focus on chromosomes 1 − 22 and X in our study.

The preprocessing starts with the following two steps: 1) we first sort the downloaded uniform peaks lexicographically by chromosome number, start position and end position; 2) the human genome is divided into non-overlapping 200 base-pair (bp) bins and sorted lexicographically. The sorting benefits while running genomic operations such as intersections with the human genome for computing labels of each peak file.

Next, we compute the overlaps of 200 bps with the narrowPeaks data. For each 690 experiment, we run the intersection operation genome-wide with all 200 bps non-overlapping bins by constraining over a minimum overlapping of 50%. Thus it gives us all the 200 bp bins that have at least one overlapping of minimum 50% with any peaks. We noticed that when the same 200 bp bins correspond to different peaks in the same TF binding dataset of a specific cell, it generates the same label for such bins. In another word, it creates the duplicate bins with the same label. We removed those cases by keeping the first. Next we merged all the labels corresponding to the same 200 bp bins. The total number of 200 bins that binds to at least one TF from 91 cell lines is 1, 903, 712. Next, we remove those bins if that contains any unknown or ambiguous nucleotide, corresponding to the letter “N”, representing other than Adenine (A), Thymine (T), Guanine (G), and Cytosine(C). We found only 43 such bins, resulting in 1, 903, 669 bins remained. This is approximately 12.6% of the whole human genome. Then we define the label vector of size 690 containing 0- and 1-s, for each 200 bp bins, having at least one TF binding event. One in the *i*-th position indicates binding to the *i*-th TF-cell pair and zero means no binding to the corresponding TF-cell pair.

We make the same dataset split following DeepSEA (22) for direct comparison. Specifically, all the TF-DNA binding 200 bp bins from chromosomes 8 and 9 are kept for the test set and chromosome 7 is kept for the validation set. The remaining chromosomes are used for training the model. In Table 1, we provide the summary of our dataset. Noticeably, 85.9% and 5.3% of the bins are kept for the model training and validation; the test set consists of 8.6%. A major benefit of chromosome-wise dataset split is to defend the data leakage between the development and test set. The model never has the access to look at the motifs of chromosome 8 and 9 directly, unless the model learns the key features to identify them from the train set.

**Table 1.** Model development and evaluation dataset summary.

| Split | Chromosome (s) | Number of<br>200 bp bins | In % |
| --- | --- | --- | --- |
| Test | 8 and 9 | 165,321 | 8.6 |
| Validation | 7 | 101,892 | 5.3 |
| Train | 1-22 and X<br>(except 7, 8 and 9) | 1,636,456 | 85.9 |
| Total |  | 1,903,669 | 100 |

Note that in all the genomic arithmetic, such as intersection, merge, shuffle, overlap and so on, we use the bedtools software (42) developed by the Quinlan laboratory. To accommodate with the future update of the ChIP-seq Uniform Peaks from ENCODE/Analysis and human reference genome (GRCh37/hg19), we release the dataset generation process and generated dataset.

Figure 2 depicts the label distribution in percentage along 690 TF binding labels. To do that, we first compute the total number of labels in each dataset split separately and then compute the value in its own split. While plotting the figure we sort the values according to the test set, as such the line for the test is smoother. The plot shows many challenges regarding the TF binding prediction problem. The dataset is imbalanced in all splits, however the imbalance shows similar trend of the label distribution. In addition 70% of test data belongs to only 257 labels out of 690. To mitigate this issue we utilize the label weights computed from the train set in the cross entropy loss computation while developing models.

**Fig. 2:**
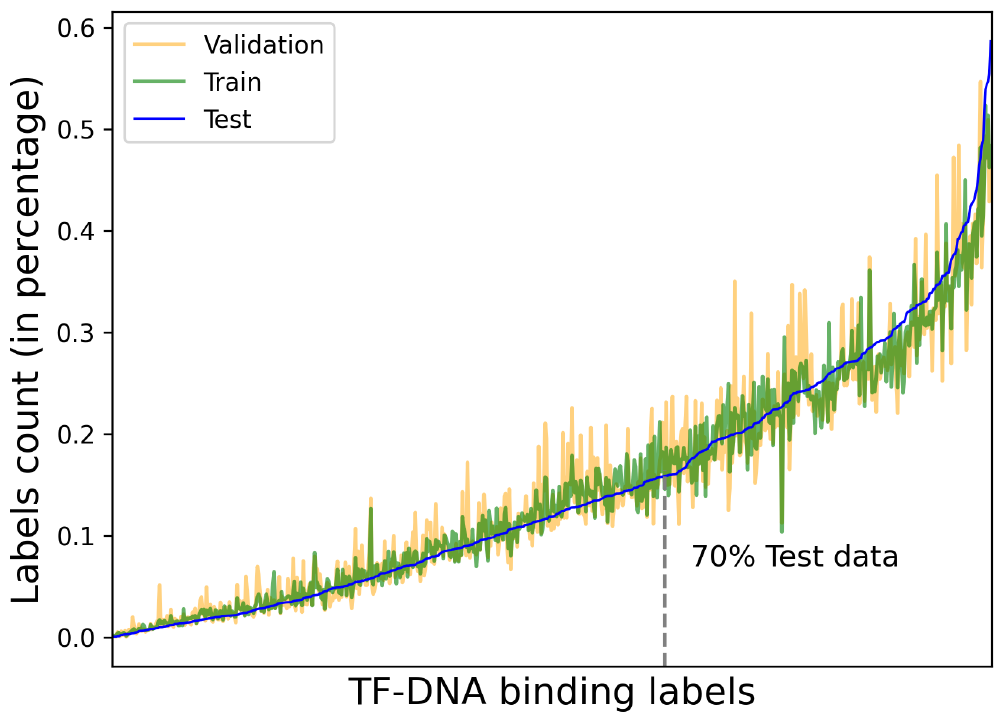
Labels distribution in ascending order percentage corresponding to each dataset split. The train, validation and test splits follow the same distribution along the labels. Only 257 labels contain 70% of the test data.

### EPBD characteristics

Given an input DNA sequence, following the Algorithm presented in (37), we run at least 100 Markov chain Monte Carlo (MCMC) simulations with different initial conditions. The temperature is set to 310.0 Kelvin with 50, 000 and 80, 000 preheating and post preheating steps, respectively, throughout the studies. Each random displacement is performed by selecting a random bp’s updated coordinate cutoff threshold of 25 °A. The breathing dynamics are recorded throughout the post preheating steps at different runs as the samples of the generated Markov chain. The distribution of the samples is computed over the independent runs and normalized over the number of post preheating steps as the final breathing characteristics.

When utilizing the pyDNA-EPBD model for analysis of short DNA sequences such as gcPBM and HT-SELEX, the addition of flanking sequences can play a critical role. This is primarily because the biophysical properties of DNA often extend beyond the confinement of the sequence of interest and are influenced by the broader bp context. The biophysical traits of a short sequence, in isolation, may not fully encapsulate these sequence-dependent effects. Thus, by including flanking sequences, we are better able to simulate the native context of the DNA sequence, enhancing the accuracy of the model’s outputs. Alexandrov *et. al* (15) revealed that the simulation distribution matches the dramatic difference in the transcriptional activity of the promoters. Following the same analysis, we compute the average coordinate distance profile (*y*_*n*_) for the all the sequences using our pyDNA-EPBD model. We also computed the average flipping profile using our pyDNA-EPBD model as the probability of a bp being flipped throughout the simulation steps. A bp is considered flipped if the coordinate distance of that bp is no less than a threshold (in °A). We collected flipping profiles at five different thresholds from 0.707106781186548 °A to 3.53553390593274 °A, inclusive, with step size 0.707106781186548. Thus, average coordinates along with five flipping profiles computed at five different thresholds comprise our DNA breathing feature matrix of dimensions feature length(200) ×6.

#### Model architecture

We have developed a multi-modal computational model specifically designed to ascertain TF-DNA binding affinities at designated chromosomal coordinates, denoted in the format chrx:y.

The core of this model revolves around a meticulously extracted DNA sequence, spanning a length *l* commencing from the position *y* on chromosome *x*. To encapsulate a comprehensive contextual understanding, we incorporate an adjoining flank of predefined size *f* . This flank, encompassing both the preceding and succeeding nucleotide bases relative to the focal point, is seamlessly integrated at both ends of the primary sequence, thereby forming an extended full context sequence of total length *L* = *f* + *l* + *f* . In our implementation, we fix the value of *l* and *f* as 200 and 400, respectively. Our model operates through a three-step process to analyze DNA sequences for transcription factor binding. Initially, it converts the DNA sequence into a tokenized format. The first key step employs a transformer encoder neural network, specially pre-trained on DNA sequences, to generate high-dimensional features representing the complex interplay of nucleotide sequences. Concurrently, the model also computes DNA breathing dynamics, focusing on the fluctuating structure of the DNA helix, which is vital for understanding transcription factor binding and gene regulation. This module creates a unique set of features representing the dynamic aspects of individual nucleotides. Finally, the feature aggregation module merges the sequence features with the DNA breathing dynamics via a sophisticated cross-attention mechanism, effectively synthesizing these distinct feature sets with different modality to highlight key interactions. The details are presented as follows:

### Module-1: Extraction of DNA high-dimensional features

This module operates on the full-context DNA sequence, which is extended to a length of *L*. The initial phase involves transforming the sequence into a tokenized form and subsequently encoding these tokens into integers, resulting in a sequence length of *L*^′^. This tokenization is a prerequisite for the subsequent computational steps. Following this, we utilize a transformer encoder neural network, which has been pre-trained exclusively on DNA sequences, including human DNA and others as shown as Module-1 in Fig. 3. The primary focus of this network’s training has been on masked-language modeling tasks, where it learns to predict the identity of masked nucleotide based on the surrounding sequence context. This training strategy enables the network to effectively capture the intricate patterns and dependencies inherent in the DNA sequence. The output of this process is a set of high-dimensional feature vectors, denoted as *x*_1_ and represented in a multi-dimensional space 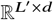 These feature vectors encapsulate the complex relationships and relative positions of nucleotides within the DNA sequence, providing a comprehensive and nuanced representation essential for accurate TF-DNA binding prediction. We utilize the DNABERT2 (43), containing 117 million parameters, as the pre-trained large language model to encode the full context DNA sequence for the subsequent modules.

**Fig. 3:**
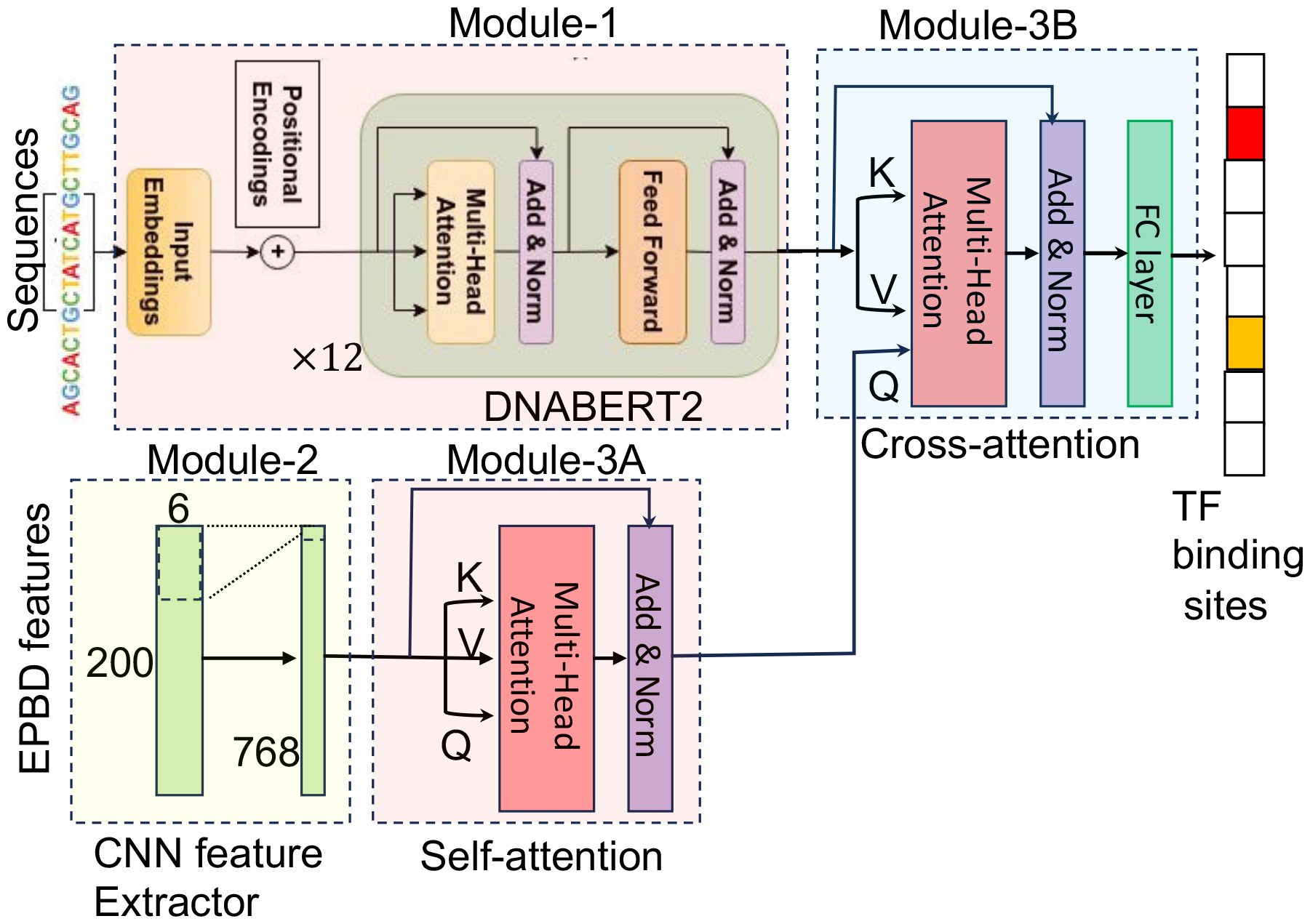
Overview of the proposed EPBD-DNABERT2 framework

### Module-2: Extraction of DNA breathing dynamics features

The second critical component of our model addresses the analysis of DNA breathing dynamics, focusing on the central DNA sequence of length *l*. DNA breathing dynamics pertain to the transient structural openings within the DNA double helix, often referred to as “DNA breathing” or “DNA bubbles.” These spontaneous structural changes are crucial for a myriad number of biological functions, including transcription, replication, and the binding of transcription factors to DNA. In contrast to the high-dimensional feature extraction previously described, which corresponds to each token of the sequence, this module is engineered to compute a distinct set of features for each nucleotide within the pivot sequence. The output of this analysis is a set of features represented as *x*_2_ ∈ ℝ^*l*×*c*^, where each feature captures distinct aspects of the DNA breathing dynamics. The variable *c* here represents the various channels or dimensions of these dynamics, encapsulating different characteristics and properties of the DNA transient structural changes.

The EPBD feature extraction module employs a one-dimensional convolutional neural network (CNN) to distill both local and distant hierarchical features from the DNA breathing dynamics as shown as Module-2 in Fig. 3. This CNN is designed to align the output dimensions with the high-dimensional feature dimensions of Module-1 (*d*), ensuring consistency in subsequent processing steps. By quantifying these dynamics at the nucleotide level, the module provides deep insights into the local conformational flexibility of the DNA. This information is crucial for understanding and predicting how transcription factors interact with specific DNA regions, as these dynamic structural changes can significantly influence TF binding affinity and specificity.

### Module-3: Multi-modal feature aggregation

The culmination of our multi-modal modeling approach lies in its feature aggregation module, which synergistically combines the independently computed high-dimensional DNA sequence features and DNA breathing dynamics. This integrative process is crucial for accurately predicting TF-DNA binding affinity. Following the EPBD feature extractor, we employ a self-attention mechanism coupled with layer normalization as shown as Module-3A in Fig. 3. This step is pivotal in identifying and emphasizing the most salient features, sized *l* × *d*, that are directly relevant to the TF-DNA binding prediction task. The ability of attention mechanism to focus on critical sequence regions enhances the model’s interpretability and prediction accuracy. Subsequently, a cross-attention module, also followed by layer normalization, effectively fuses the high-dimensional sequence features; 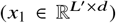 with the processed DNA breathing features 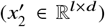as shown as Module-3B in Fig. 3. In this operation, *x*_1_ serves as the key and value, while 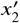 acts as the query, facilitating a comprehensive integration of the two feature sets. Further refinement of these aggregated features is achieved through a positionwise feedforward network. Comprising two fully connected layers with a ReLU non-linear activation function after the initial layer and followed by layer normalization, this network generates the final multi-modal features (*x*_3_ ∈ ℝ^*l*×*d*^). The model then applies a mean pooling layer to consolidate these features into a cohesive output, followed by another fully connected layer with ReLU non-linearity. This step computes fixed-size features representing the entire input DNA sequence context. Finally, a classification layer transitions from these fixed-sized features to the number of TF binding classes, accompanied by a sigmoid activation layer. As the prediction task is multi-label in nature, with each TF binding event being mutually exclusive, the sigmoid activation yields the probability of the input DNA sequence binding to each TF in the set.

### Hyperparameters optimization

In our quest to refine the multi-modal framework for TF binding site prediction, a critical aspect was the hyperparameter optimization comprising more then 500 models. The model’s complexity and the intricate nature of TF-DNA interactions necessitated a fine-tuned approach to identify the optimal set of hyperparameters that maximize predictive accuracy. The hyperparameter optimization was conducted using the Ray Tune framework, coupled with the Optuna and Hyperband hyperparameter optimization library. We employed a comprehensive search strategy focusing on various model parameters, including the number of heads in the multi-head attention mechanism, kernel size of the EPBD embedder, dropout rate, learning rate, and weight decay. These hyperparameters were crucial in adjusting the model’s capacity to learn and generalize from the dataset comprising DNA sequences and their corresponding epigenetic markers. The search space for the optimization process was defined as follows:

- **Weight Decay**: Loguniform distribution from 1*e* − 4 to 1*e* − 1.
- **Learning Rate**: Loguniform distribution from 1*e* − 6 to 1*e* − 2.
- **EPBD Embedder Kernel Size**: Discrete choices of 3, 5, 7, 9, and 11.
- **Dropout Probability (p dropout)**: Uniform distribution from 0.01 to 0.5.
- **Number of Attention Heads (num heads)**: Discrete choices of 2, 4, 8, 16, and 32.

Our optimization process involved training the model across various combinations of the aforementioned hyperparameters. The training and validation datasets were constructed using the multi-model framework, encompassing DNA sequences and their epigenetic characteristics. Model training was executed for a predefined number of epochs, with batch sizes and data loading workers set to optimize computational efficiency. The Ray Tune framework facilitated the efficient exploration of the parameter space, while the Optuna library’s pruning strategies expedited the search by dismissing sub-optimal trials. The primary metrics for model performance evaluation were validation loss and Area Under the Receiver Operating Characteristic Curve (AUROC), with a focus on minimizing the loss and maximizing the AUROC.

The outcome of this extensive hyperparameter optimization process was a robust set of parameters that significantly enhanced the model’s ability to accurately predict TF-DNA binding sites. The best configuration, as determined by the lowest validation loss, encompassed an optimal combination of weight decay of 0.000125, learning rate of 1.216*e* − 5, EPBD embedder kernel size of 7, dropout probability of 0.0195, and the number of attention heads of 2. This hyperparameter combination provided test loss of 0.0481 and overall test AUROC of 0.950 This configuration was instrumental in achieving superior predictive performance, as evidenced by the improvement in AUROC and Area Under the Precision-Recall Curve (AUPR) metrics.

#### Other benchmark models

These models are developed as part as this study, consequently another contribution for benchmarking the performance of the final proposed model. Note that, we apply imbalanced class weights computed from the train set in the loss function while training any model.

### Finetuned DNABERT2

This model is a variant of the original DNABERT2, fine-tuned to adapt to the specific nuances of our dataset. It serves as a baseline to measure the incremental benefits of incorporating EPBD features.

### DNABERT2 with random EPBD feature aggregation

This model variant integrates EPBD features with DNABERT2 representations, but the aggregation is random. This helps to assess the impact of structured versus unstructured integration of EPBD features.

### DNABERT2 with vanilla EPBD feature aggregation

In contrast to the random aggregation model, this version employs a more systematic approach to integrating EPBD features with DNABERT2. This model is essential to demonstrate the effectiveness of our feature aggregation strategy.

## Results and Discussions

Breathing dynamics improve human TF-DNA binding sites prediction. We thoroughly investigated pyDNA-EPBD features both *in vivo* and *in vitro* experimental datasets. While applying coordinate and flipping features in isolation *in vitro*, we also examine the applicability of the breathing characteristics in the multi-modal context with a genomic pretrained large language model. We report the improved performance and study our underlying physics-enabled system in the human TF-DNA binding specificity prediction both in classification and regression tasks. In the following sections, we discuss the TF-DNA binding sites prediction in the ChIP-seq experiments and, subsequently in the gcPBM and SELEX datasets.

### Breathing dynamics (BDs) improve *in vivo* human TF-DNA binding prediction

Using a physically constructed stability function as a guide, we trained and evaluated the resulting EPBD features in a multi-modal application. We report on the area under the receiver operating curve (AUROC) and area under the precision-recall curve (AUPR) metrics for assessment and comparison with the state-of-the-art models. To do so, we first apply the model to the test set, which is derived from the chromosome 8 and 9 (which together account for more than 8.6% of the human genome). The model is asked to assess whether each 1000-bp sequence from the test set binds to a TF (antibody) in a specific cell environment in a mutually exclusive manner. This results in AUROC and AUPR scores along 690 the TF-DNA binding events. Next we report the average AUROC and AUPR across all TF-DNA binding events in Table 2. We report the performance of TBiNet (27), DanQ (1), and DeepSEA (22) as baseline models, as described by the TBiNet. DanQ and TBiNet solve the problem as a multi-label classification tasks as the same setting of ours. As part of this study, the DNABERT2-finetuned model—an additional baseline and contribution—is adjusted for the downstream tasks of TF-DNA binding prediction. Overall our model performs on par with the TBiNet, however it outperforms others specifically DNABERT2-finetuned model by a 3% relative increments on average in both AUROC and AUPR metrics. The improvement relative to the DanQ is also significant, particularly in AUPR metric, more that 3%. Improvements in both metrics demonstrate that the breathing dynamics aid the DNABERT2 model to increase the true-positive rates and reduce the false-positive rates that essentially improves the TF-DNA binding events prediction tasks.

**Table 2.** Overall performance comparison of our model (EPBD-DNABERT2-crossattn) with baseline works in light of AUROC and AUPR metrics. EPBD-DNABERT2-crossattn performs on par with TBiNet and outperforms others, specifically DanQ and DNABERT2-finetuned, by a 3% relative margin on AUPR metric.

| Methods | AUROC | AUPR |
| --- | --- | --- |
| DeepSEA | 0.901 | 0.248 |
| DanQ | 0.931 | 0.295 |
| TBiNet | 0.947 | 0.333 |
| **Ours** |  |  |
| DNABERT2-finetuned | 0.918 | 0.296 |
| EPBD-DNABERT2-crossattn | 0.949 | 0.326 |

Next, we analyze and study genome-wide TF binding sites with respect to cell environment according to the same performance metrics. EPBD-DNABERT2-crossattn improves over 665 TF-DNA binding tasks out of 690 when comparing only AUROC metric, 655 tasks while only considering AUPR compared with DNABERT2-finetuned model. Figure 4 depicts all cases using AUROC and AUPR metrics separately. The diagonal line determines which model performs best, specifically lower-right and upper-left triangle shows the space for DNABERT2-finetuned and EPBD-DNABERT2-crossattn, respectively. Our model scales up to 9.6 percent in the AUROC metric. The Figure clearly demonstrates that in most TF-DNA binding tasks the AUROC metric scales up from 0.85 which determines the model’s true capacity to recognize the TF-DNA binding sites from the physics guided genome information. Since the class distribution is highly imbalanced, the baseline thresholds for each TF-DNA binding tasks are different in AUPR metric. Thereby showing relative improvement of 3% compared to the baseline DNABERT2-finetuned establishes the quantitative performance boost. The bottom panel additionally showcases the EPBD-DNABERT2-crossattn’s superiority compared to the DNABERT2-finetuned model. Figure 4 also shows that the model down-performs mostly in the binding factor when considering cell line K562, which is the lymphoblast cells used in immune system disorder and immunology research. From the deep look in the dataset we find that there are 150 TF binding events in cell line K562, and EPBD-DNABERT2-crossattn performs less in 9 of them only. In addition, we find that our model suffers at most in the TF binding of BDP1 antibody in cell type K562 and HeLa-S3.

**Fig. 4:**
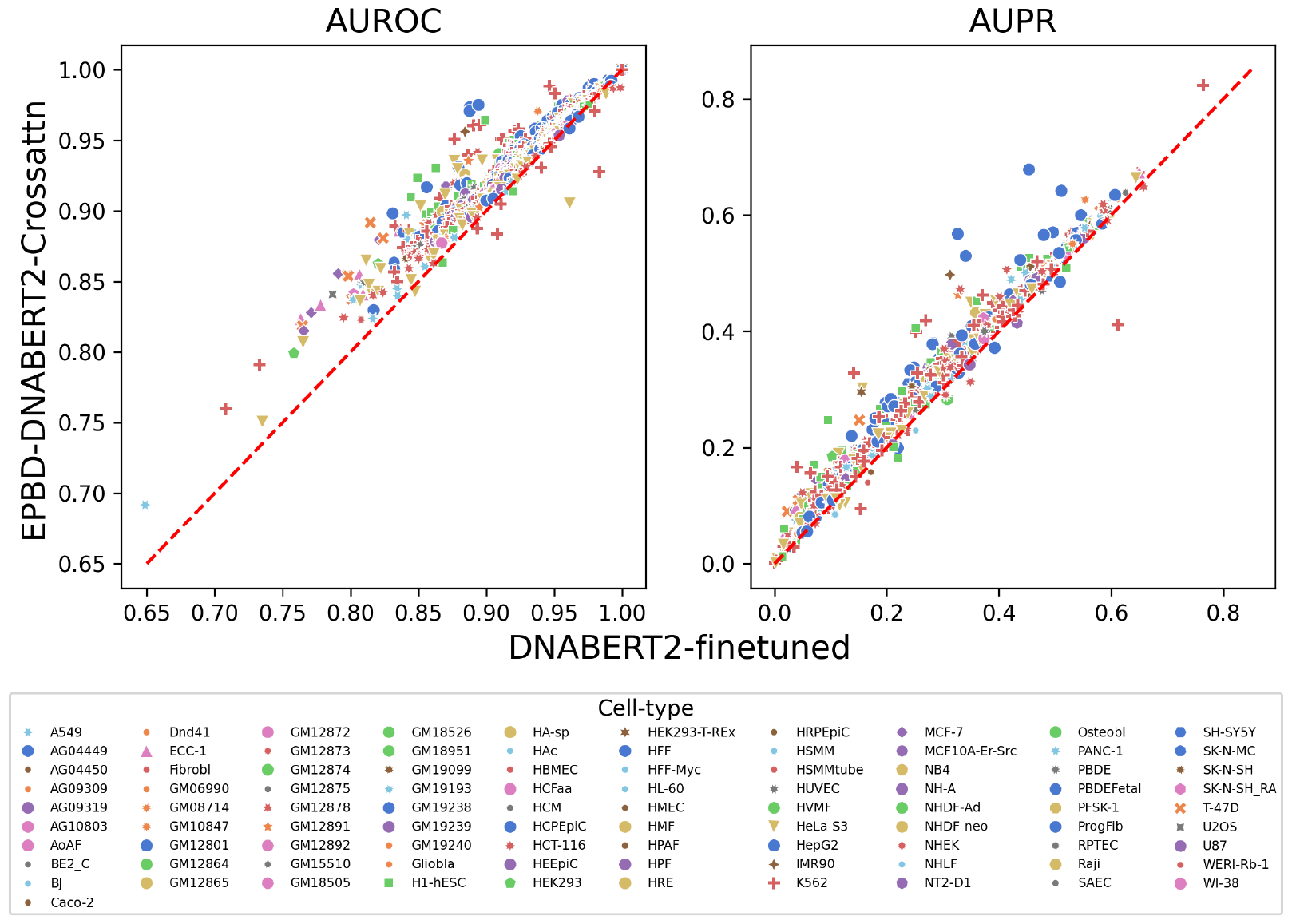
Relative performance improvement of EPBD-DNABERT2-crossattn over DNABERT2-finetuned in AUROC (left panel) and AUPR (right panel) metrics. The scores are colored by the cell line information which demonstrates that breathing dynamics guides DNABERT2 to improve on both metrics across different cell types.

To understand the top improvements EPBD-DNABERT2-crossattn over DNABERT2-finetuned, we plot the top 20 relative improvements using false positive rate versus true positive rate. Figure 5 shows that the combination of cell type and antibody where the improvements are most significant. The bold and dotted lines correspond to the EPBD-DNABERT2-crossattn and DNABERT2-finetuned performance, respectively, with same color span denoting the cell type and antibody combination. The improvements scale from 7.1% to 9.6%. The Figure clearly demonstrates that our model significantly improves the TF-DNA binding sites prediction across different antibodies and cell types, particularly MafF and MafK.

**Fig. 5:**
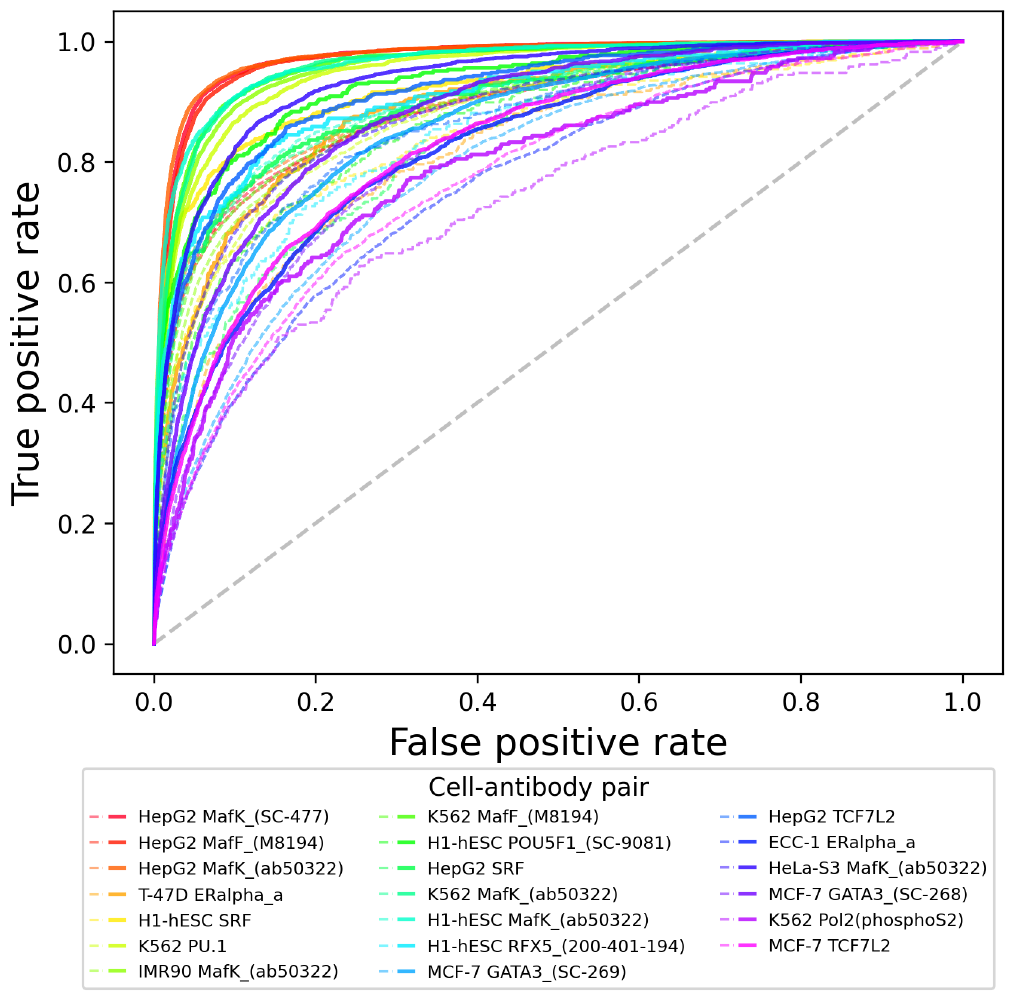
Top 20 relative improvements of EPBD-DNABERT2-crossattn over DNABERT2-finetuned model. Right panel provides the corresponding label information (cell type and antibody pair) by bold and dotted lines denoting EPBD-DNABERT2-crossattn and DNABERT2-finetuned, respectively. The contribution of EPBD features is diverse across different cell types and antibodies.

### EPBD features guides large language model

To enhance the interpretability of the cross-attention mechanism within our EPBD-DNABERT2 model, we performed an in-depth analysis of the singular value spectrum derived from cross-attention weights. For a diverse set of 10, 000 input sequences, we compared the influence of actual EPBD features against randomly generated features on the model’s decision-making process. The singular values extracted from the cross-attention matrix—when actual EPBD features were utilized—demonstrated a significant and consistent divergence from those obtained using random features, indicating the meaningful contribution of EPBD features in guiding the model towards more accurate TF-DNA binding predictions. This was visually captured in Figure 6 where two distinct curves represent the normalized cumulative singular values, with the EPBD feature curve markedly elevated, reflecting the EPBD features’ fine-grained impact in directing the model’s predictive functionality.

**Fig. 6:**
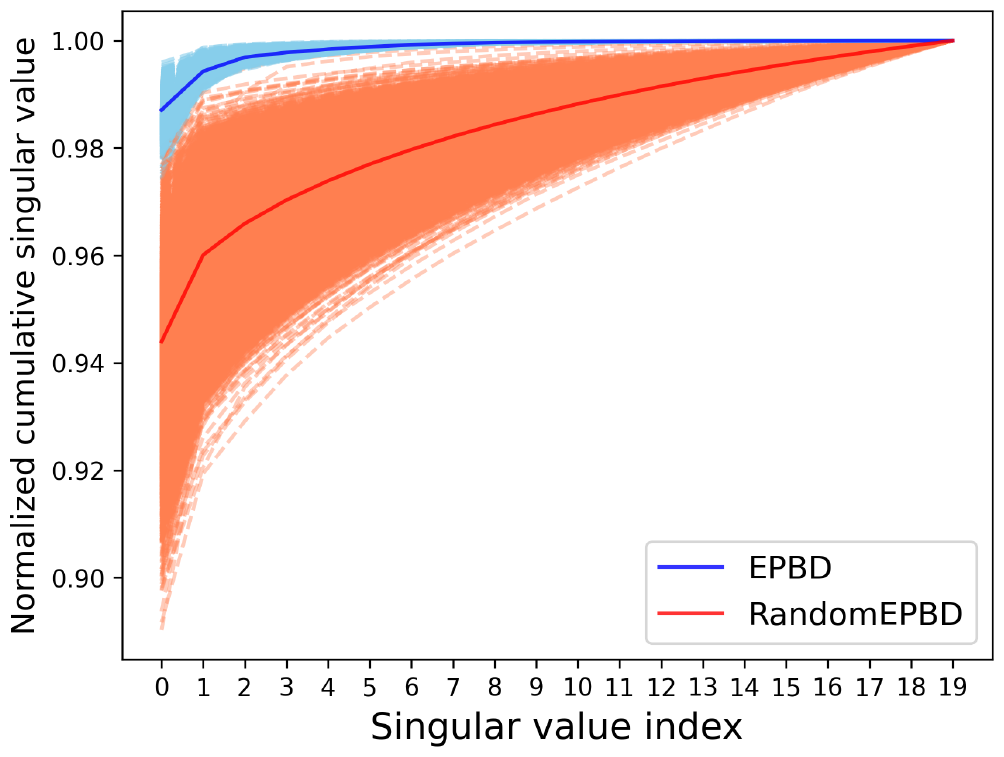
Normalized cumulative singular values of 10, 000 input DNA sequences corresponding to EPBD-DNABERT2-crossattn (blue) and randomly permuted EPBD features (red) shows that EPBD breathing dynamics guide DNABERT2 for extracting meaningful information.

Our comprehensive analysis of EPBD-DNABERT2 model’s cross attention mechanism included a focused evaluation of the top 20 singular values from the cross-attention weights for 10,000 input sequences. These values were normalized and cumulatively plotted in Figure 6, showcasing the variance when using actual EPBD features versus randomly permuted EPBD features. The Singular values act as indicators of key abstract features crucial for prediction, with the EPBD features employed as queries and DNABERT2 features as key and values in the aggregation process. This approach underscores the substantial role of EPBD features in refining the model’s decisions, evidenced by the distinct divergence in singular value spectra between EPBD and random features.The gap highlights EPBD features effectiveness in enhancing model precision, contrasting with the baseline performance of random features. Notably, the EPBD-DNABERT2-crossattn model displays a more nuanced expressiveness, with the cumulative values starting on average of above 0.98 when leveraging pivotal singular values, compared to 0.95 for random features. This analysis demonstrates the impact of EPBD features in guiding the DNABERT2 model, akin to a physics guided system, for genome wide TF-DNA binding event prediction.

The cross-attention heatmaps reveal significant differences in the model’s performance with EPBD features for different transcription factors as shown in Figure 7. For the improved performance scenario, there is a pronounced focus in the center of the heatmap, suggesting that the inclusion of EPBD features sharpens the model’s attention on essential sequence regions, thereby enhancing TF prediction accuracy. This focused attention likely corresponds to the model picking up on critical sequence motifs or structural features indicative of TF binding sites. In other words, the heatmap corresponding to the scenarios with enhanced TF prediction performance showcases a concentrated attention pattern, indicating that EPBD features effectively guide the model to focus on critical regions within the sequence. This targeted attention is likely due to the model leveraging EPBD features to discern key biophysical properties of DNA, aligning with regions where transcription factor binding is most probable. Conversely, the heatmap indicating degraded performance presents a more dispersed pattern of attention, implying a less effective concentration on relevant sequence elements when EPBD features are present. This diffusion of attention could be due to the EPBD features introducing complexity or noise, which hinders the model’s ability to consistently pinpoint the salient features across the sequences, resulting in a decline in predictive reliability. The marked contrast between the two heatmaps underscores the influence of EPBD features, which can either aid in honing the model’s predictive capabilities or introduce confounding factors that obscure crucial sequence information.

**Fig. 7:**
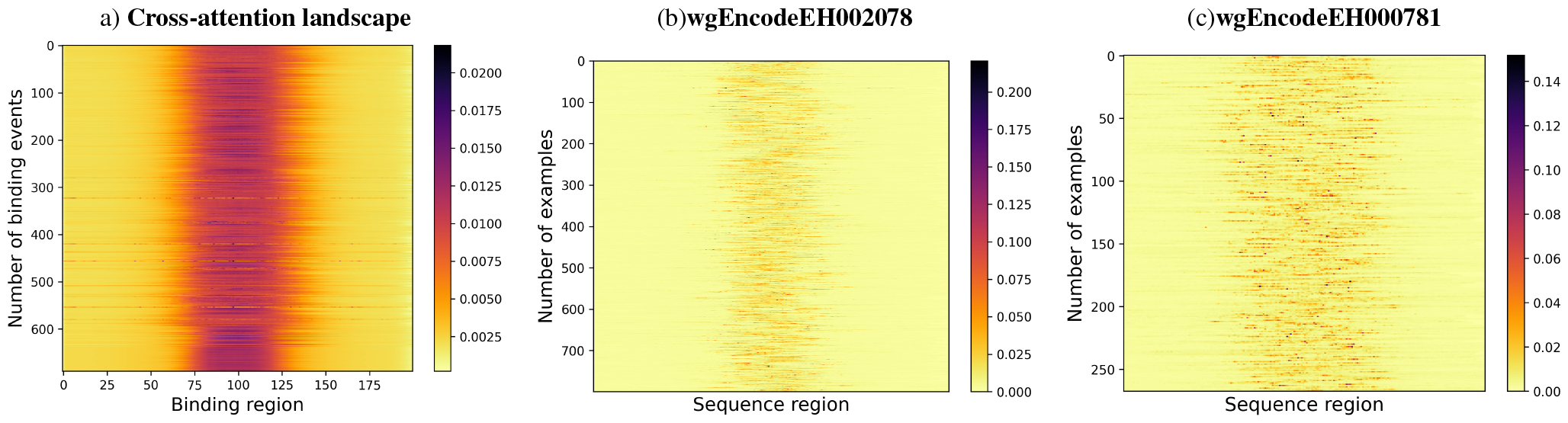
Cross attention landscape analysis of test dataset. Average cross attention landscapes for (a) all 690 TF-dna binding events, (b) wgEncodeEH002078 as of good quality and (c) wgEncodeEH000781 as of caution quality TF-DNA binding experiments are depicted. The model is able to discriminate the binding region from the non-binding region across many input DNA sequences.

The additional analysis of the EPBD CNN feature extractor weights, as presented in Figure 8, offers further insights into the model’s internal feature prioritization mechanisms. The kernel weight distribution across the EPBD feature channels elucidates the underlying significance of each feature in influencing the model’s predictive capabilities. The plot illustrates the mean and median values for the weight distribution across six channels. The channels associated with a feature extractors that utilizes coordinate-based EPBD features and flipping profiles delineated by five distinct thresholds, as detailed in Subsection 2.1.3. This arrangement suggests that each channel processes specific biophysical aspects of DNA, with one dedicated to spatial coordinates and the remainder to dynamic flipping events occurring at varying threshold levels. The line graph in Figure 8 clearly shows that channels 3 and 5 exhibit the most significant fluctuations in weight values, with channel 3 displaying the highest mean weight and channel 5 showing a notable dip below zero before recovering. This suggests that the feature extractor within the EPBD CNN is attributing the highest importance to the features processed in channel 3, as indicated by the positive peak, implying a direct correlation with the target variables. In contrast, the negative peak for channel 5 might represent an inverse relationship, where the features in this channel contribute negatively to the model’s output. Channels 1, 2, 4, and 6 display less volatility in weight values, indicating a more consistent but less pronounced impact on the model’s predictions. The median values follow a similar trend to the means, underscoring the central role of these channels in the feature processing hierarchy. The weight distribution, particularly the prominence of channels 3 and 5, is crucial for understanding how the CNN feature extractor prioritizes EPBD features before passing them to subsequent self-attention and cross-attention mechanisms for TF binding prediction.

**Fig. 8:**
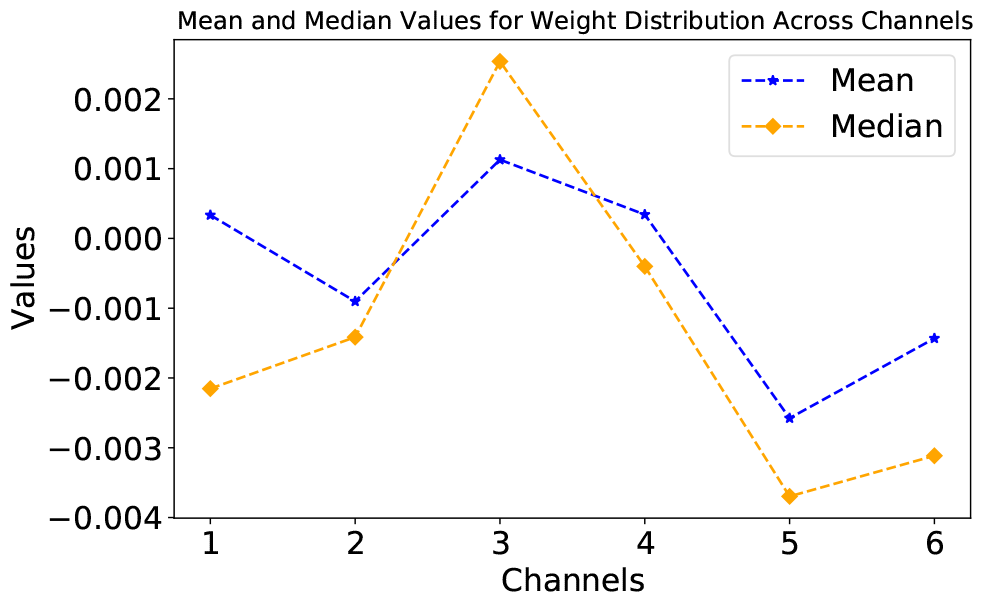
Distribution of EPBD feature extractor weights.

### Motif Discovery

The EPBD-DNABERT2 model demonstrates proficiency in extracting conserved motifs from extensive DNA sequences, employing a methodology akin to the one delineated for DNABERT (30). In contrast to DNABERT, which harnesses all self-attention heads, our model capitalizes solely on cross-attention values to assess the influence of pyDNA-EPBD features. Given a DNA sequence encompassing 1000 base pairs, our analysis is contingent on a cross-attention matrix dimensioned at [200 × *l* ], stemming from the tokenized *l* tokens paired with the central 200 nucleotide bases’ breathing dynamics. This design allows for a broader contextual awareness of both binding and non-binding regions. To distill this information into a more nucleotide-centric perspective, we calculate the average cross-attention weights along the *l* tokens, thus yielding attention scores for individual tokens that are modulated by EPBD features. The normalization of these attention values relative to the nucleotide count within each token results in an average cross-attention weight at the single nucleotide resolution. These weights are instrumental in pinpointing regions abundant in motifs, which we subsequently corroborate against the JASPAR CORE (2022) (44) non-redundant experimental motif database, utilizing the TOMTOM (45) motif comparison tool for validation purposes.

In our quest to delineate motifs within DNA sequences, we initiate by identifying regions of heightened attention using average cross-attention weights, leveraging a protocol inspired by DNABERT. To ascertain a contiguous high attention region, we set forth three criteria: first, the cross-attention value for each base must exceed the sequence’s average; second, it must be at least twentyfold the sequence’s minimum value—a stringent threshold relative to DNABERT’s tenfold to account for the heightened specificity attributed to EPBD features; and third, the region’s length must be twice that considered by DNABERT, pinpointing longer motifs with a length of ten. These regions of high cross-attention are preliminarily earmarked as motifs. A subsequent hypergeometric test refines this selection, retaining only those motifs meeting an adjusted p-value threshold of 0.005, in alignment with DNABERT’s standard. Given that each sequence in the test set is marked by a TF-DNA binding site, these are presumed to be motif-rich, forming our positive set, while an equal number of generic non-binding sequences constitute our negative set. We used P-values, e-values, and q-values as statistical metrics to assess the significance of motifs, which are distinctive patterns within biological sequences such as DNA, RNA, or proteins. A p-value calculates the probability of a specific motif occurring by random chance, with a lower p-value indicating greater statistical significance. The e-value extends this concept by considering the number of motif searches conducted, predicting the number of times a motif might appear by chance in a given dataset; hence, a lower e-value strengthens the case for the motif’s biological relevance. The q-value adjusts the p-value to account for the false discovery rate when multiple comparisons are made, with a lower q-value indicating a reduced likelihood of falsely identifying a motif as significant. These values collectively provide a robust framework for identifying and validating biologically meaningful motifs in sequence data.

Motifs passing this statistical scrutiny are deemed high-quality candidates. A pairwise alignment among all such motif pairs is then conducted, merging similar motifs when the alignment score surpasses the greater of the two benchmarks: either one less than the required contiguous region length or half the length of the shorter motif in the pair. To transpose these into a PWM-like format, we standardize the length by centering and extracting a fixed-length window around each identified motif, selecting a length of 15 to approximate the average motif width reported in JASPAR CORE (2022). Final filtration steps include discarding motifs with fewer than ten instances, ensuring that only the most robust motifs are retained. This meticulous process culminated in the discovery of 124 high-quality motifs, a subset of which are depicted in the referenced Figure 9, differentiated by their e-value—a measure of quality. Further details and correlations to known motifs are expounded upon in the accompanying table 3, underscoring the model’s adeptness in recognizing motif-rich regions across diverse motif families.

**Table 3.**
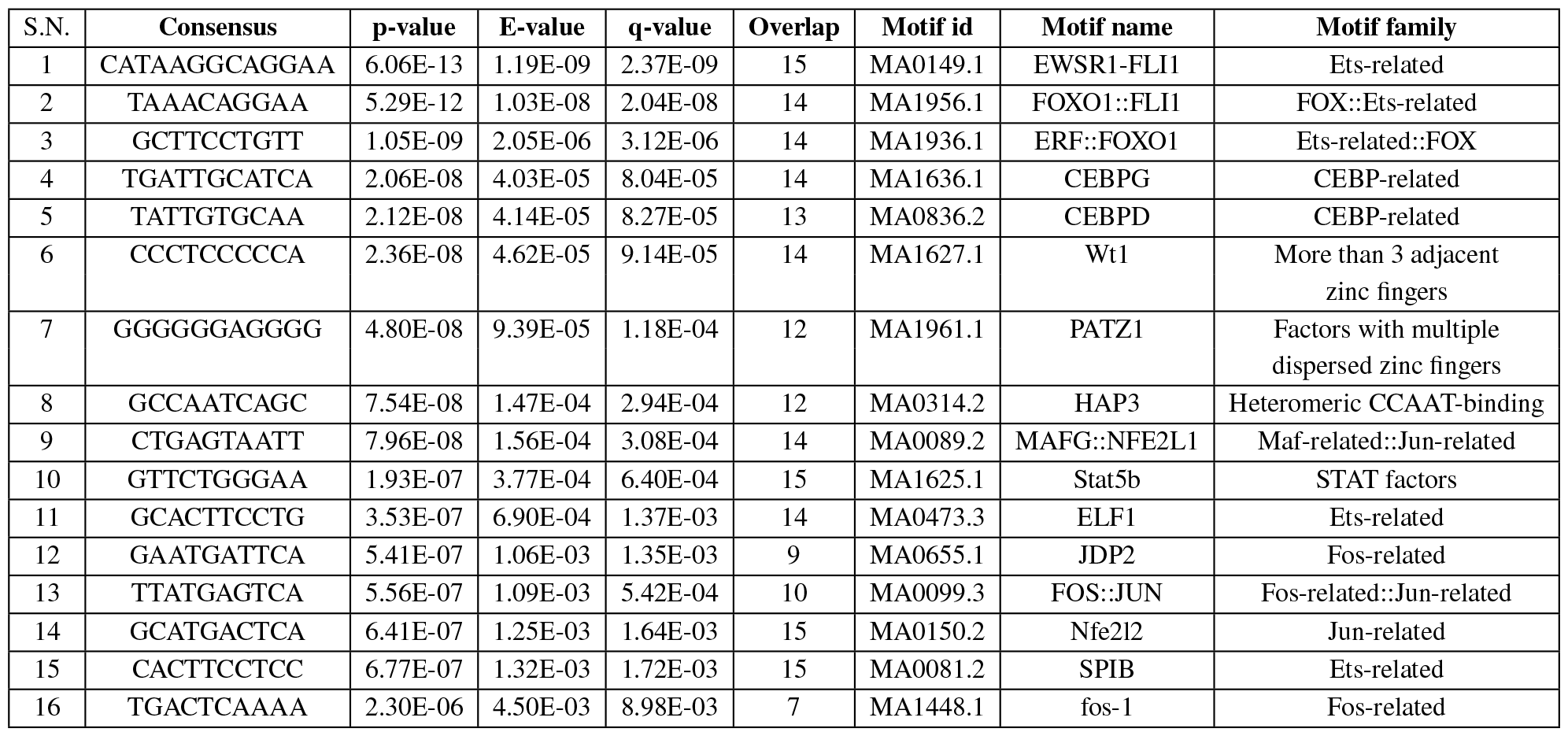
Selected motifs from multi-modal framework analysis with cross attention weights. The motifs are sorted in ascending order by the e-value column. The motif id column corresponds to JASPAR2022 (44) database.

**Fig. 9:**
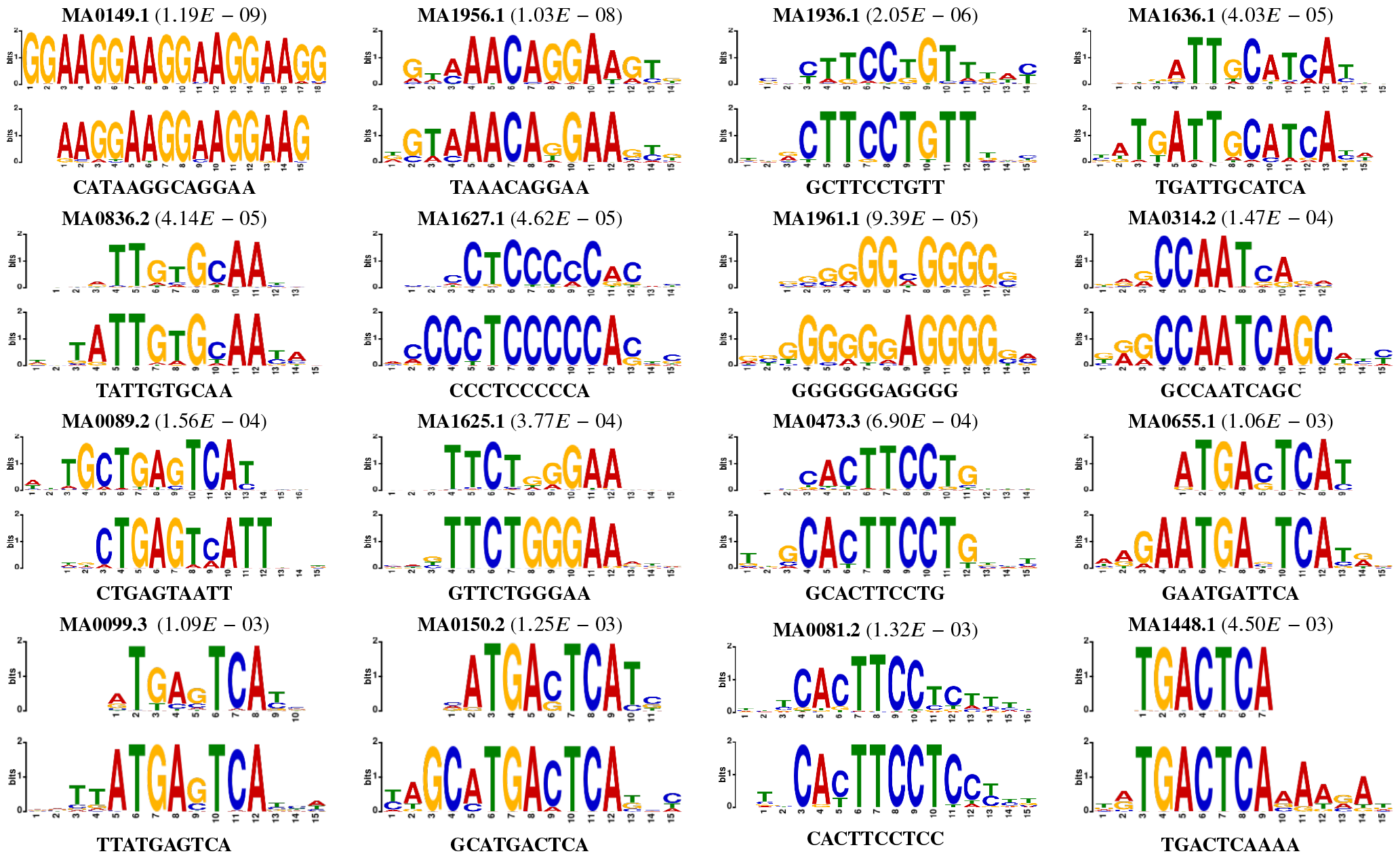
Visual motif comparison using weblogos corresponding to motifs related to Table 3. The top panel shows the motif reported by the JASPAR2022 (44) database, and the bottom panel is the corresponding motifs found by EPBD-DNABERT2-crossattn in each panel.

Based on the results, the analysis yielded a diverse collection of motifs across various transcription factor families and classes, underscoring the model’s capability to discern a broad spectrum of DNA-binding preferences. The statistical rigor of the findings is substantiated by low p-values and E-values, denoting a high degree of significance and a low likelihood of these motifs arising by chance. Notably, motifs linked to the TEAD3, FOS::JUN, and STAT1 transcription factors stand out due to their biological relevance in gene regulatory networks. The E-values, particularly for motifs like CEBPG and MAFG, suggest that these motifs are not just statistically significant, but also biologically interesting, pointing towards their potential regulatory importance. The overlap with known motifs from JASPAR further validates the model’s predictive accuracy and its utility as a tool for biological discovery. In particular, the motifs related to EWSR1-FLI1, manifesting the lowest E-values, underscore the model’s precision in pinpointing motifs of exceptional interest. Such motifs, characterized by their rarity and specificity, are likely to be pivotal in the regulatory landscapes they inhabit. The recovered motifs correlate with known TF classes, such as bZIP and Ets-related factors, which are instrumental in gene expression regulation. Overall, the table presents compelling evidence that the integration of cross-attention mechanisms and biophysical features within a deep learning framework can effectively pinpoint crucial motifs that may have substantial implications for understanding the complexities of gene regulation. It is worth noting that the discovery of motifs provides us with an interpretation the model’s findings.

### Utility of breathing dynamics (BDs) in *in vitro* TF-DNA binding datasets

In this study, we extend the application of EPBD features to evaluate their utility in quantifying transcription factor (TF)-DNA binding specificity. Utilizing the biophysical attributes of DNA, specifically the dynamics of nucleotide flipping and spatial coordinates, we conducted a regression analysis on *in vitro* TF-DNA binding specificity datasets from gcPBM (19) and SELEX (20**?** ). Given the concise sequence lengths and sparse sample sizes, we opted for a Support Vector Regression (SVR) model with a radial-basis function kernel, renowned for its effectiveness with high-throughput quantitative datasets, over a transformer-based approach. Two distinct approaches were considered for the analysis: firstly, the sequences were encoded using one-hot encoding, representing the standard sequence information alone; secondly, we incorporated the breathing characteristics of the sequences, which includes both coordinate and flipping information, to assess the added value of these dynamic features. The comparison of these two approaches was carried out through a 10-fold cross-validation procedure, enabling us to measure and contrast the predictive performances when sequences are analyzed both with and without the integration of their associated breathing dynamics. This methodical evaluation aims to ascertain the extent to which the inclusion of DNA’s biophysical dynamics can enhance the accuracy of TF-DNA binding specificity predictions.

Table 4 presents the comparative results of a 10-fold cross-validation analysis for predicting TF-DNA binding specificities, focusing on Mad, Max, and MYC transcription factors within the gcPBM dataset. The gcPBM dataset comprises 23, 028 sequences, each originally 36 base pairs (bps) in length, which were extended to 44 bps by flanking each side with *GCGC*. For our analysis, only the central 36 bps—exclusive of the flanks—were used to consider the breathing dynamics. Our findings indicate a minimum of 3% enhancement in the coefficient of determination (*R*^2^) when breathing dynamics are considered alongside the sequence information, as opposed to relying on sequence information alone. Remarkably, the features generated by the pyDNA-EPBD model account for an average of 91% of the variance in the observed data. Moreover, the reduced standard deviation in cross-validation scores suggests a consistent performance improvement when breathing dynamics are included, implying a more reliable prediction model compared to using sequence data alone.The negative mean-squared error (MSE) metric further underscores the predictive improvement, revealing at least a 3% enhancement when breathing dynamics are integrated. The negative MSE, being closer to zero, denotes a more accurate prediction of binding affinities.

**Table 4.** Performance of sequence alone versus sequences with pyDNA-EPBD breathing dynamics (coordinate and flip) with SVR-RBF. This reports average *R*^2^ and negative mean-squared error (MSE) performance metrics of 10-fold cross validation with standard deviation.

| TF | Sequence<br>$R^2$ | Sequence<br>MSE(-ve) | Sequence+BDs<br>$R^2$ | Sequence+BDs<br>MSE(-ve) |
| --- | --- | --- | --- | --- |
| Mad | 0.880±0.005 | -0.120±0.004 | 0.931±0.003 | -0.069±0.003 |
| Max | 0.890±0.004 | -0.110±0.004 | 0.920±0.002 | -0.080±0.003 |
| Myc | 0.861±0.006 | -0.139±0.005 | 0.897±0.004 | -0.103±0.004 |

In summary, the incorporation of breathing dynamics, as characterized by the pyDNA-EPBD model, has shown substantial value in refining the prediction of TF-DNA binding affinities, thereby evidencing the significant predictive power of DNA’s biophysical properties in computational binding affinity models.

Our regression analysis was further extended to 215 SELEX TFs spanning 27 distinct families. The SELEX dataset is extensive, with over 19 million base pairs distributed across approximately 1.78 million sequences. These sequences vary in length from 9 to 16 base pairs. To facilitate the simulation, we standardized the sequences by flanking each with a consistent sequence pattern *GCGCGCGCGCGCGCGCGCGCGCGCGC*, elongating them to a minimum of 61 and a maximum of 66 base pairs.

Figure 10 presents a visual comparison of the regression scores, plotting the coefficient of determination (*R*^2^) for sequences alone (x-axis) against those supplemented with breathing dynamics (y-axis). The placement of nearly all data points above the diagonal baseline in the Figure indicates that the integration of breathing dynamics consistently outperforms the sequence-only approach in predicting TF-DNA binding specificity. This consistent position above the baseline highlights the added predictive value that breathing dynamics bring to the table, suggesting that the biophysical properties encoded by these dynamics are not just incidental but are indeed influential in determining TF-DNA interactions. The enhanced predictive accuracy afforded by including breathing dynamics underscores their potential as a valuable asset in computational models for large-scale analysis of TF-DNA binding specificity.

**Fig. 10:**
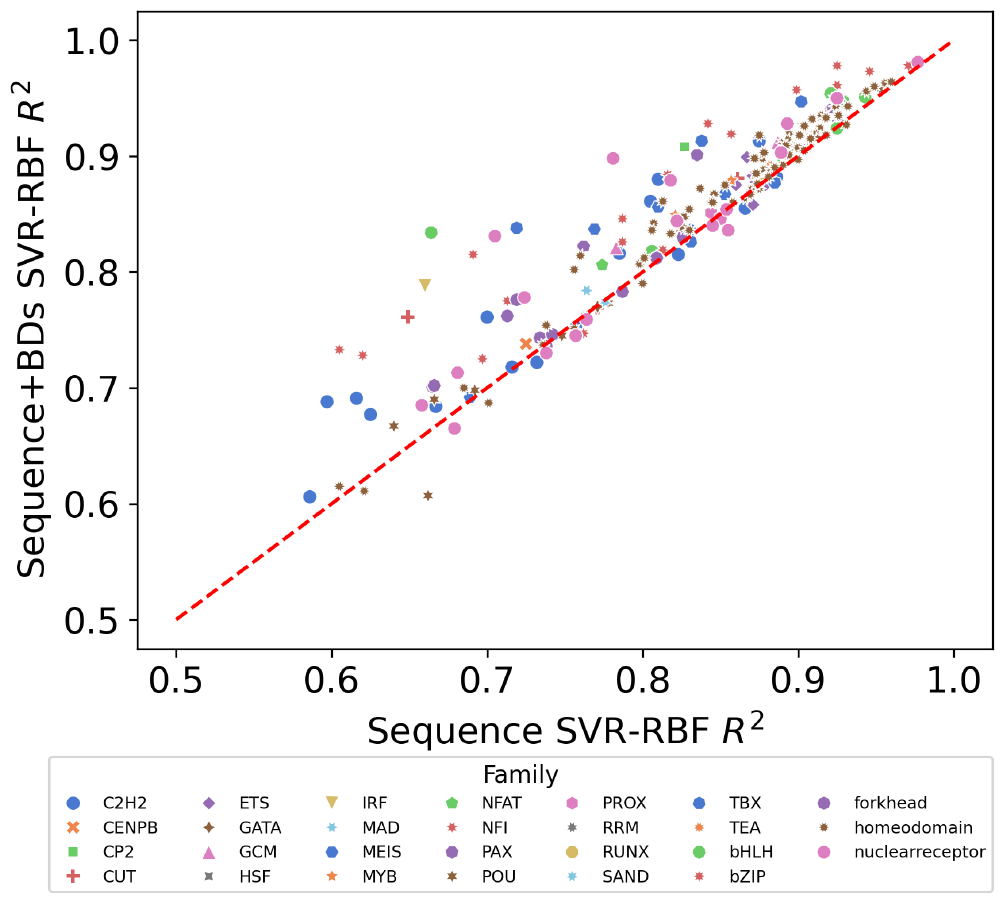
Average *R*^2^ of 10-fold cross validations using SVR-RBF on SELEX TF-binding scores. The Figure shows useful utility of breathing dynamics over sequence alone information.

### Ablation Study

The ablation study delineated herein provides a systematic examination of the constituent elements of the EPBD-DNABERT2 framework. By methodically removing or altering specific components, we aim to distill the individual contributions of these components to the model’s overall performance in transcription factor-DNA binding prediction.

### Cross-attention vs. Simple Multi-modal Input Features Aggregation

An integral part of our analysis focuses on the cross-attention mechanism and its comparative efficacy against simpler strategies for the aggregation of multi-modal input features. Such elementary strategies include the direct concatenation or summation of input features. Through this comparison, we aim to validate the hypothesis that cross-attention provides a superior method for the nuanced integration of EPBD features with the pre-trained nucleotide representations of DNABERT2.

### Coordinate vs. Flip vs. Coordinate-plus-flip Features

Further dissecting the feature space, we investigate the individual and combined impact of coordinate and flip features derived from EPBD. Each feature type encapsulates distinct aspects of the biophysical properties of DNA. By evaluating these features in isolation as well as in conjunction, we seek to understand their respective and cumulative influence on the model’s predictive accuracy. This examination will elucidate whether a synergistic effect is observed when both feature types are utilized in tandem.

In the results presented in Table 5, we notice a marked improvement in the performance metrics when cross-attention is employed, as evidenced by the highest scores in both AUROC and AUPR. This underscores the efficacy of the cross-attention mechanism in capturing the intricate dependencies between EPBD features and nucleotide sequences, leading to enhanced predictive performance.

**Table 5.** The ablation study shows the performance of different model configurations with respect to the AUROC and AUPR metrics. Our final proposed multi-modal EPBD-DNABERT2-crossattn model has at least 2% relative improvement on both scales.

| Methods | AUROC | AUPR |
| --- | --- | --- |
| DNABERT2-finetuned | 0.918 | 0.296 |
| RandEPBD-DNABERT2 | 0.917 | 0.296 |
| VanillaEPBD-DNABERT2-coord | 0.921 | 0.311 |
| VanillaEPBD-DNABERT2-coordflip | 0.920 | 0.308 |
| EPBD-DNABERT2-crossattn | 0.949 | 0.326 |

## Conclusion

In conclusion, our comprehensive study on the EPBD-DNABERT2 model has demonstrated its robustness and efficacy in predicting transcription factor (TF)-DNA binding sites. By integrating biophysical DNA properties, particularly EPBD features, with the advanced DNABERT2 foundation model through a cross-attention mechanism, we’ve achieved notable improvements in binding site prediction accuracy. The study highlights the importance of a multi-modal approach, blending computational and biophysical insights, thereby offering a novel perspective in understanding genomic interactions. Our ablation studies underscored the significance of cross-attention in feature integration and the complementary roles of different EPBD feature types. These findings not only enhance our understanding of TF-DNA interactions but also open new avenues for exploring genomic data, with potential implications in biomedical research and therapeutic development. Our analysis on motif discovery additionally provides with a clear, sequence-level interpretation of the model’s findings. The versatility of our model, evidenced by its performance across diverse datasets, signifies a substantial step forward in computational genomics, paving the way for more nuanced and effective analysis of complex biological mechanisms.

## Supporting information

justification

## Declarations

### Availability of data and materials

EPBD-BERT is Open Source Software published under the 3-Clause BSD License and can be found at https://github.com/lanl/EPBD-BERT.

### Authors’ contributions

AK and MB developed and implemented EPBD-BERT, and AK performed all validation experiments. MB, KOR, ARB, AU, AS, and BSA consulted AK for analysis and validation of results. All authors wrote and edit the manuscript.

## Footnotes

1 http://rohslab.cmb.usc.edu/MSB2017/

2 http://hgdownload.soe.ucsc.edu/goldenpath/hg19/encodeDCC/wgEncodeAwgTfbsUniform/

3 https://hgdownload.soe.ucsc.edu/goldenPath/hg19/bigZips/latest/

