## Supplementary material for "Advancing Transcription Factor Binding Site Prediction Using DNA Breathing Dynamics and Sequence Transformers via Cross Attention": justification

In the updated version, we have incorporated references pertaining to HT-SELEX and gcPBM datasets. Furthermore, we have included and elaborated on the foundational papers that introduced the concept of DNA shape features. These papers highlight the significant improvement in Transcription Factor (TF) binding specificity predictions when both sequence and shape models are utilized, compared to the use of sequence data in isolation. The added references are follow:


```
@article{htselex-and-gcpbm-data,
  title = {Expanding the repertoire of DNA shape features for genome-scale studies of transcription factor binding},
  volume = {45},
  ISSN = {1362-4962},
  url = {http://dx.doi.org/10.1093/nar/gkx1145},
  DOI = {10.1093/nar/gkx1145},
  number = {22},
  journal = {Nucleic Acids Research},
  publisher = {Oxford University Press (OUP)},
  author = {Li,  Jinsen and Sagendorf,  Jared M. and Chiu,  Tsu-Pei and Pasi,  Marco and Perez, Alberto and Rohs,  Remo},
  year = {2017},
  month = nov,
  pages = {12877–12887}
}

@article{zhou2015quantitative,
  title={Quantitative modeling of transcription factor binding specificities using DNA shape},
  author={Zhou, Tianyin and Shen, Ning and Yang, Lin and Abe, Namiko and Horton, John and Mann, Richard S and Bussemaker, Harmen J and Gord{\^a}n, Raluca and Rohs, Remo},
  journal={Proceedings of the National Academy of Sciences},
  volume={112},
  number={15},
  pages={4654--4659},
  year={2015},
  publisher={National Acad Sciences}
}
@article{yang2017transcription,
  title={Transcription factor family-specific DNA shape readout revealed by quantitative specificity models},
  author={Yang, Lin and Orenstein, Yaron and Jolma, Arttu and Yin, Yimeng and Taipale, Jussi and Shamir, Ron and Rohs, Remo},
  journal={Molecular systems biology},
  volume={13},
  number={2},
```

pages={910},
year={2017}
}

@article{zhou2013dnashape,
title={DNAshape: a method for the high-throughput prediction of DNA structural features on a genomic scale},
author={Zhou, Tianyin and Yang, Lin and Lu, Yan and Dror, Iris and Dantas Machado, Ana Carolina and Ghane, Tahereh and Di Felice, Rosa and Rohs, Remo},
journal={Nucleic acids research},
volume={41},
number={W1},
pages={W56--W62},
year={2013},
publisher={Oxford University Press}
}
